# A minimal-complexity light-sheet microscope maps network activity in 3D neuronal systems

**DOI:** 10.1101/2022.06.20.496852

**Authors:** Paulina M Wysmolek, Filippo D Kiessler, Katja A Salbaum, Elijah R Shelton, Selina M Sonntag, Friedhelm Serwane

## Abstract

*In vitro* systems mimicking brain regions, brain organoids, are revolutionizing the neuroscience field. However, characterization of their electrical activity has remained a challenge as this requires readout at millisecond timescale in 3D at single-neuron resolution. While custom-built microscopes used with genetically encoded sensors are now opening this door, a full 3D characterization of organoid neural activity has not been performed yet, limited by the combined complexity of the optical and the biological system. Here, we introduce an accessible minimalistic light-sheet microscope to the neuroscience community. Designed as an add-on to a standard inverted microscope it can be assembled within one day. In contrast to existing simplistic setups, our platform is suited to record volumetric calcium traces. We successfully extracted 4D calcium traces at high temporal resolution by using a lightweight piezo stage to allow for 5Hz volumetric scanning combined with a processing pipeline for true 3D neuronal trace segmentation. As a proof of principle, we created a 3D connectivity map of a stem cell derived neuron spheroid by imaging its activity. Our fast, low complexity setup empowers researchers to study the formation of neuronal networks *in vitro* for fundamental and neurodegeneration research.

## 1. Introduction

Electrophysiological readouts at single-neuron resolution are one building block to our understanding of the brain. A major hurdle, however, is the limited experimental access to living human brain tissue granted by state-of-the-art recording technologies. In recent years, tremendous progress has been made with the introduction of cerebral organoids, multicellular systems derived from embryonic and induced pluripotent stem cells^1-3^. These *in vitro* models recapitulate aspects of *in vivo* neuron behavior and tissue architecture. To what extent they replicate network function as seen *in vivo* is an open question^1,4^.

As the first functional studies of 3D neuronal networks emerge^4^, the experimental challenges remain strict: Neurons generate and propagate electrical signals at a millisecond timescale via action potentials. Thus, the functionality of a neuronal tissue needs to be assessed at this timescale in 3D, all at a single-neuron resolution. Patch clamps record electrophysiological signals of single neurons within neurospheres^5^, cerebral organoids^6-8^ and retinal organoids^9-11^ with sub-millisecond resolution. However, only a few individual neurons can be inspected at once which prevents the mapping of larger networks, a signature of the brain. Multi-electrode arrays (MEA) capture extracellular potentials created by neuronal populations. The technology has been used to analyze 2D connectivity and neural circuits in neurospheres,^12^ as well as in brain and retinal organoids^3,13-16^. Recently, initial 3D characterization of network behavior in brain organoids was performed by combining MEA with shank probes^4^. While electrode approaches offer unprecedented time resolution, their spatial (z) resolution remains constrained by the ability to place electrodes at various depths. In addition, information about which neuron types build up the network remains hidden due to a lack of access to image cell morphologies. This is a particular drawback for characterizing organoids consisting of different neuron types, for example cerebral^15^ and retina^17,18^ organoids. In those systems, it is hypothesized that neuron circuits play the same crucial role for signal processing as in their *in vivo* counterparts^1,19,20^. Monitoring 3D network activity with simultaneous access to cell morphology and cell type is possible via calcium^21^ and voltage^22^ imaging using fluorescence microscopy. The functionality of both cerebral and retinal organoids has been characterized with one- and two-photon confocal microscopy^17,23-25^. However, experiments have been limited to 2D due to the (point) scanning speed of confocal microscopes. To open this speed bottleneck, custom microscopes have been developed in order to parallelize the acquisition. Sculpted light approaches use the temporal properties of light pulses to allow the simultaneous illumination of different depths^26,27^, at the cost of a complex pulsed laser setup. Lightfield microscopes are significantly simpler setups which record both position and angle information of emitted rays using a microlens array. While this allows fast computational sectioning of individual planes, the axial resolution remains limited^28^. This has been overcome in parts by training a neuronal network with images recorded with a light-sheet microscope^29^. Lightsheet microscopy provides, due to the planar illumination profile, true optical sectioning and a plane-wise parallel readout, typically limited only by the framerate of the camera^30^. Functional activity mapping in various samples has been performed at scanning rates between 0.5 to 50 Hz (Suppl. Table 1)^31-37^. High-speed imaging using a single primary objective both for illumination and detection has become possible via oblique plane microscopy, introduced by C. Dunsby^38^. While these setups are in principal suited to perform volumetric imaging of neuronal networks, they require advanced optical expertise in their assembly and alignment. Efforts to reduce their complexity rely on the integration of excitation and illumination optics in one module (OCPI)^39^. Although this simplifies the optical setup, the mechanical setup remains complex as it requires special sample mounting and moving the module at a few millisecond intervals. Approaches to further reduce complexity introduced lightsheet modules as addon to standard inverted microscopes or minimalistic standalones^40,41^. While add-ons exist which provide fast volumetric recording at sub-cellular resolution^42,43^, they require external optical paths for illumination, or a confocal microscope as a basis (Leica SP8/Stellaris DLS). Consequently, volumetric Ca-imaging with 3D single neuron resolution on a minimalistic platform still is an unsolved problem.

Here, we introduce a minimalistic custom-built light-sheet microscope tailored to state-of-the-art computational neuroscience tools^44^. The innovation of our approach is (i) a minimalistic module replacing the condenser of a standard inverted microscope together with standard glass bottom dishes as sample mounts (ii) fast volumetric imaging by a custom lightweight piezo stage which has the footprint of a 96-well plate (iii) a processing pipeline which reduces typical lightsheet artifacts such that state of the art neuroscience tools can be used for downstream processing. Our setup allows for calcium imaging of a range of samples in physiological conditions with subcellular resolution. At the same time, it operates at a speed comparable with complex custom-built microscopes. It is tailored to widely used calcium-imaging probes, but displays orders of magnitude lower complexity and costs (<50 k€) compared to previous implementations (Suppl. Table 1). As a proof of principle, we used our platform to record 3D neuronal network activity and created a global connectivity map of the network within three-dimensional neuron cultures. While our results constitute the first step in the activity mapping of 3D neural cultures, the platform is versatile and can be used with a range of samples from neuro-spheres to mature organoids.

## 2. Experimental setup and analysis pipeline

High speed light-sheet illumination integrated into widely-used off the shelf microscopes and standard glass bottom dishes would serve a broad community of researchers. We implemented a plug and play light-sheet module as an add-on to a standard inverted microscope where it simply replaces the microscope condenser (Fig. 1). The lightsheet is aligned via the screws typically used for adjusting Köhler illumination, making the alignment easily accessible to non-experts. To reduce complexity of the optics to a minimum, a static planar light-sheet configuration generated by a cylindrical lens L0 and a spherical symmetrical lens L1 was chosen and mounted via machined opto-mechanics components. To allow for the generation of thinner lightsheets, e.g. for cell imaging, an aspheric lens was chosen for L1. This enables diffraction limited focusing with an illumination NA=0.3 when a larger focal length for L2 is chosen. For a flexible choice of fluorescent reporters and alignment-free light delivery, a commercial laser combiner was used. The setup contains three fiber-coupled laser lines at 488 nm, 561 nm and 643 nm. Light intensity was set via the laser combiner’s analog inputs using an Arduino controlled via Micromanager. For Ca imaging applications an observation volume of 666×666×60 μm^*3*^ (333×333×60 μm^*3*^) was recorded with a voxel size of 1.3×1.3×3 μm^3^ (0.65×0.65×3 μm^3^) using the 20x (40x) objective. As the diameter of typical organoid samples exceed 1 mm, a specific subsection is recorded.

**Figure 1:**
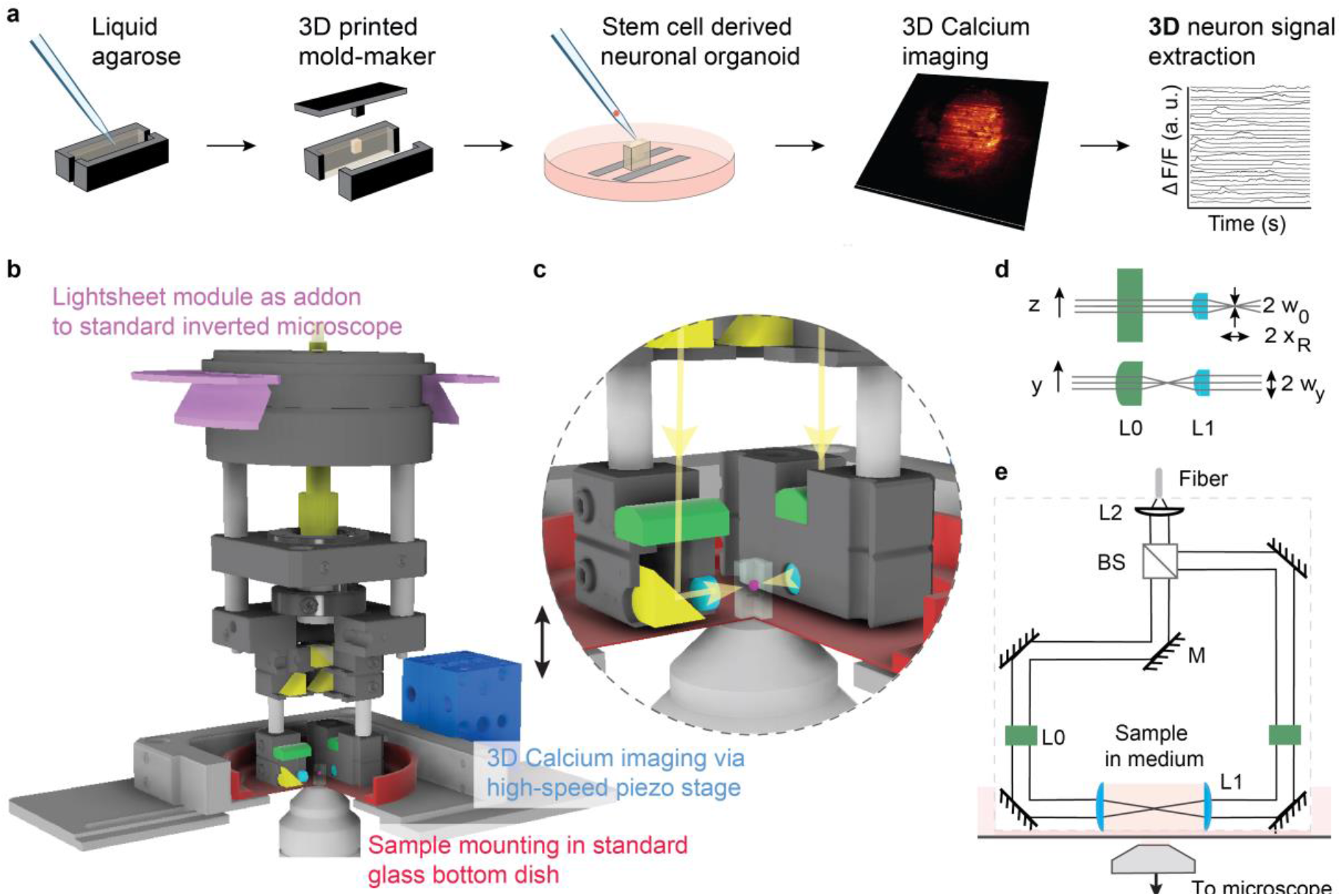
A minimal complexity light-sheet microscope enables fast volumetric imaging of neuronal signals. **(a) Workflow for the extraction of 3D neuron signals**. Temporal segmentation of neuron traces becomes possible via a custom pre-processing pipeline in combination with recent 3D neuron analysis software^44^. **(b,c) The light-sheet module as an add-on to a standard inverted microscope**. Volumetric imaging at single neuron resolution is achieved by movement of the sample along the optical axis using a high-speed piezo-stage (10 ms per plane or 5 volumes/s with 20 planes/volume). **(d,e) Light-sheet optical path**. The optical beam, delivered via a fiber, is split via a cube beam splitter and directed to the sample by right-angle prisms (M). A static planar light-sheet is created by focusing the beam via a cylindrical lens (L0) at the back focal plane of an aspherical lens (L1).

### 2.1 Flexible sample mounting and imaging

To allow mounting of a variety of samples and placing the light-sheet module in their close proximity, we used standard 50mm glass bottom dishes (Willco Wells, HBST-5040). We minimized sample movements by embedding it in an agarose well (Fig. 1a). To image deep into the tissue long working distance water dipping objectives (20x: Olympus UMPLFLN20XW; 40x: UMPLFLN40XW) were used (Fig. 1e). While the objectives are designed to operate without cover glass, we reasoned that a cover glass together with water as immersion medium will not impact their resolution because of their long working distance combined with their moderate NAs (20x: wd = 3.5 mm, NA = 0.5; 40x: wd = 3.3 mm, NA = 0.8). We verified that the resolution is independent of penetration depths by imaging beads embedded in 1% agarose (Suppl. Fig. S4). To characterize chromatic shifts in illumination and detection, we imaged multicolor beads which we excited using all three wavelengths. We find that axial and lateral chromatic shifts remain on the order of the optical resolution, thus not substantially impacting imaging at several wavelengths (Suppl. Fig. S4b). Physiological conditions (37°C, 5% CO_2_) were maintained by enclosing the microscope with a custom-built incubation chamber made out of Plexi glass.

### 2.2 Acquisition speed tailored for Ca-imaging

To resolve responses of typical genetically encoded calcium indicators (GECIs), tens-or hundreds millisecond temporal resolution per volume is needed. To acquire volumetric scans the sample is moved through the light-sheet step-wise via a custom piezo-stage (Fig. 1b). We maximized the volumetric scanning rate by minimizing the load on the stage and inverting the direction of movement after each z-stack completion. As a result, the setup allowed for volumetric scanning at 10 ms/plane, currently limited by the rate of camera acquisition (100 fps). In contrast to other simplistic setups^39^, the sample is moved while the illumination and detection optics remain still. Due to the lower weight of the glass bottom dish compared to the objective, this, in principle, allows faster scanning.

### 2.3 Subcellular optical resolution

To determine the microscope’s resolution, we characterized the light-sheet parameters and the system’s point spread function (PSF) (Fig. 2). The lateral 1/e resolution mainly depends on the numerical aperture of the objective. The axial resolution is predominantly influenced by the thickness of the anisotropic Gaussian beam light-sheet in case of the 20x objective due to its finite optical sectioning capabilities (NA=0.5). The Gaussian beam is parametrized by its beam waist *w*_0_ which determines its thickness as 2*w*_0_ and its Rayleigh length *x*_*R*_ = *n π w*_0_^2^/*λ* with wavelength *λ* and refractive index *n*. Its lateral extension is given by its 1/*e*^2^ width *w*_*y*_. To quantify those parameters, we imaged the light-sheet’s transversal illumination profile by deflecting the light into the detection path using a 45° tilted mirror. Different planes along the propagation axis *x* were imaged by translating the mirror with respect to the light-sheet and the objective. Each image was fitted with a 2D Gaussian and *w*(*x*) and *w*_*y*_(*x*) was extracted. Fitting *w*(*x*) yields the light-sheet’s waist *w*_0_ and its Rayleigh length *x*_*R*_ (Fig. 2a-c). The characterization was performed for all three laser lines. The thickness of the light-sheet remains well below 10 microns and thus is comparable to typical light-sheet setups (Suppl. Table 1). The recorded chromatic focus shift is on the same order of magnitude as the Rayleigh length of the light-sheet beam, allowing simultaneous multicolor imaging (Fig 2b) at decreased axial resolution (Suppl. Fig. S4). The light sheet dimensions are about a factor of 1.5 larger than the theoretical estimation. We suspect imperfection on the cylindrical lens as cause as its surface roughness is not specified to < λ/4. To measure the 3D resolution, we recorded the PSF by imaging gel-embedded 170 nm fluorescent beads (Fig. 2d). For each bead, we measured the lateral and the axial intensity profiles along two perpendicular axes in three central planes (*xy, xz, yz*) and fitted a Gaussian function. The 1/*e* radius *σ*_*x,y,z*_ was taken as the resolution measure yielding a lateral resolution of 0.9 µm (0.5 µm) and an axial resolution of 1.7 µm (1.1 µm) for the 20x (40x) objective, respectively. With both objectives, single-neuron resolution in all 3 dimensions is obtained in agreement with the theoretical prediction (Fig. 2e).

**Figure 2:**
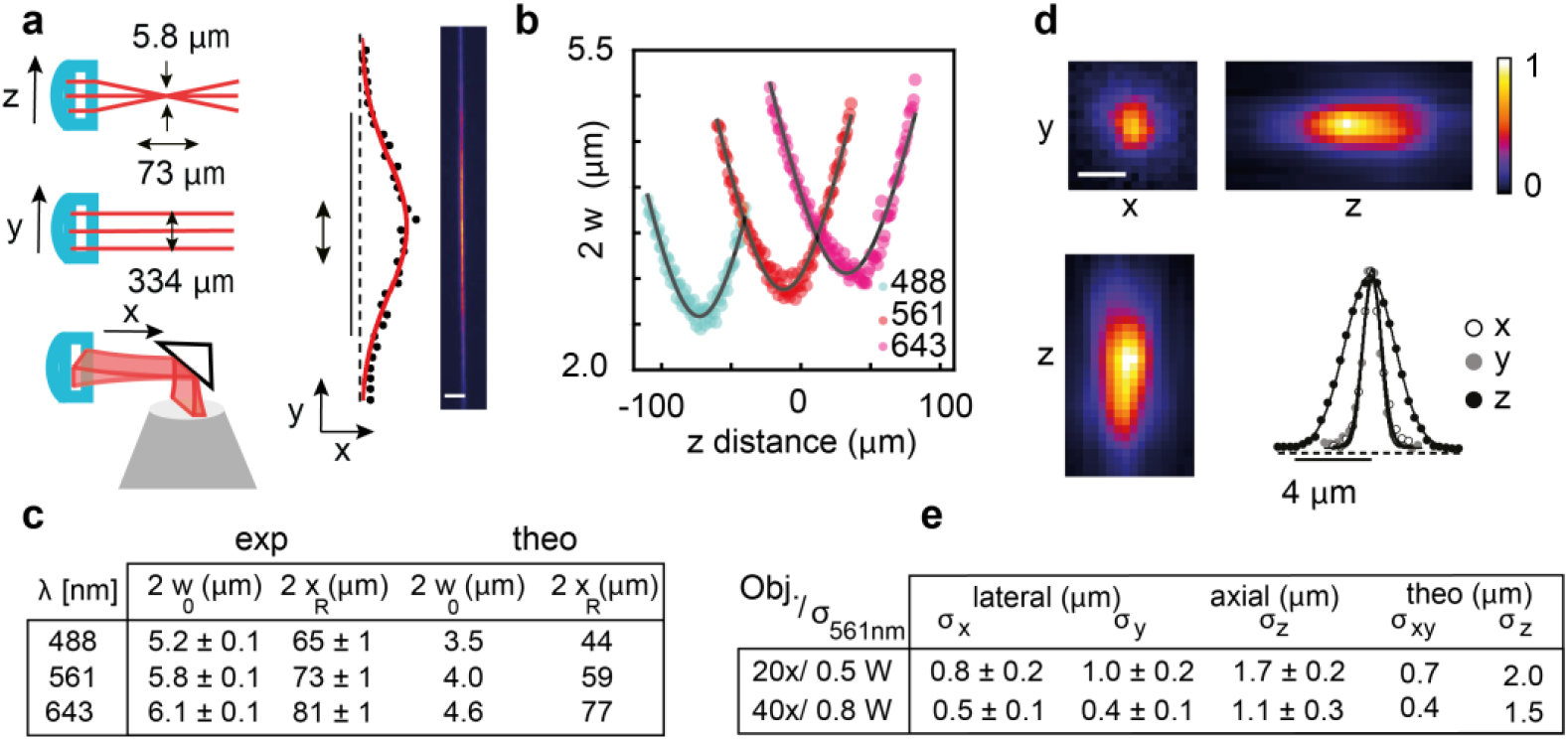
A simplistic light-sheet microscope with sub-cellular resolution. **(a-c) Characterization of the light-sheet parameters**. **(a)** The light-sheet propagates along the x-axis as an elliptical Gaussian beam characterized by its thickness 2 w_0_, its Rayleigh length x_R_ and its width 2 w_y_. The parameters were determined by imaging the transversal illumination profiles. **(b)** Parameters w_0_ and x_R_ are quantified by fitting a hyperbolic function (black line) to the measurements w(z). The light-sheet thickness remains well below 10 microns, allowing optical sectioning of the sample. Multicolour imaging is possible as the Rayleigh length is comparable to the chromatic shift **(c). (d-e) Quantification of the 3D resolution**. The PSF was measured by imaging gel-embedded fluorescent beads. **(e)** Values displayed in the table are mean 1/e radius (σ_x,y,z_) for profiles along the three axes with standard deviation measured for multiple beads for two objective lenses using the 561 nm laser. Scale bars: 25 μm **(a)**, 1 μm **(d)**. Number of measurements n=6 (20x; **c**,**e**); n=12 (40x,xy; **e**), n=7 (40x,z;**e**).

### 2.4 Culture of stem cell derived 3D neuronal networks

3D neuron cultures were grown according to retina organoid protocols^45^ using mouse embryonic stem cells (mESCs) (see Methods). To record calcium activity, a mESC line endogenously expressing a red fluorescence calcium sensor (R-GECO1.0) (Fig. 3, Suppl. Fig. S2) was used. To allow imaging of the positions of individual cells, the cell line constitutively expressed a nuclear marker, a *H2B-GFP* fusion protein. First, embryoid bodies were formed via the aggregation of a pre-defined initial number of mESCs cells in U-bottom well plates (3000 cells per well). Neural differentiation was triggered and proceeded for the next 7 days via the addition of Matrigel and differentiation medium under physiological conditions (5% CO2, 37°C). To optimize differentiation into specific neuron subtypes, the sample was then transferred to 40% O2, 5% CO2 and 37°C^45^. While the maximum sample growth time was limited to two weeks (see Methods), imaging and characterization was performed on day 12. The neural identity of the 3D culture could be confirmed via immunofluorescence staining (Fig. 3, Suppl. Fig. S2).

**Figure 3:**
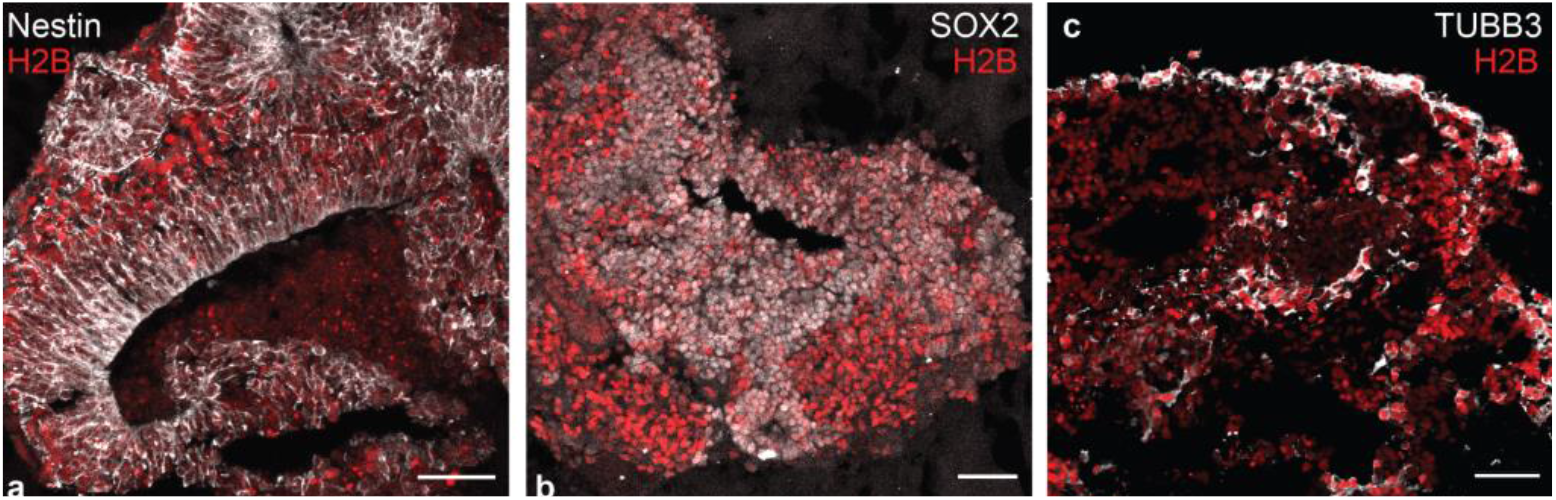
Confocal microscopy images of cryo-sections of the stem cell derived neuron culture. The presence of Nestin **(a)** and SOX2 **(b)** within rosette-like structures are signatures for neural progenitors (SOX2) and neuroepithelial cells (Nestin), respectively^64-67^. Both cell-specific markers are present in neurosphere cultures^68,69^ as well as retinal neurospheres^70^. **(c)** Class 3 ß-tubulin (TUBB3) binds microtubules in neurons and is also present in neurospheres^69,71^. Scale bar: 50 μm.

### 2.5 Optimized imaging parameters

To image neurons with sufficient axial resolution, planes were spaced at less than half a neuron diameter, Δ_Z_ = 3 µm. Furthermore, the volumetric acquisition speed was matched to the typical decay time of fluorescent Ca sensors. The R-GECO sensor has a 0.09 ± 0.02 s time-to-half-rise and 0.78 ± 0.13 s time-to-half-decay of action potential-evoked fluorescence^46^. As higher temporal resolution increases the performance of the algorithm we oversampled the decay time by a factor of 4. The scanning time in our setup can be increased for brighter indicators with faster dynamics. To reduce the size of acquired data while preserving sufficient spatial resolution, we performed 4×4 binning in *xy* to obtain recording volumes with 512 × 512 × 20 voxel, leading to a voxel size of (1.3×1.3×3) μm^3^ for the 20x objective and (0.65×0.65×3) μm^3^ for the 40x objective. With these settings, we achieve a volumetric rate of 5 Hz within an observation volume of (666×666×60) μm^3^ and (333×333×60) μm^3^, respectively, which is comparable to state-of-the-art setups (Suppl. Table 1).

### 2.6 Neuron trace extraction

Extracting the activity of densely packed neurons requires (i) the separation of spatially overlapping neurons and (ii) their discrimination from the background signal containing non-firing neurons. While several analysis packages are available (SIMA^47^, CNMF-E^48^, Suite2p^49^, ABLE^50^, SCALPEL^51^, MIN1PIPE^48^, SamuROI^52^), we chose CaImAn^44^ because of its ability to operate on 3D data^44^. First, motion correction is performed via image registration. Next, the detection of regions of interest (ROI) is carried out followed by the extraction of the temporal fluorescence signal and static neuron segmentation using a constrained non-negative matrix factorization (CNMF). To distinguish active neurons from the fluorescent background, the 4D recordings are separated into three main contributions: the spatiotemporal Ca2+ activity *AC*, a background term *B* accounting for long term drifts and out of focus fluorescence and a noise term *E*. In a final step, the reconstructed signal is obtained as

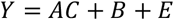

To minimize the large-scale fluorescence background not caused by Ca-fluctuations we introduced crucial pre- and postprocessing steps as highlighted in Fig. 4. First, to correct for sample bleaching, we normalized the pixel intensities of each frame by its median. After this, two light-sheet-typical artifacts remained^30^: First horizontal stripes appeared from the shade of optically dense objects. Other lightsheet setups minimize this effect by beam scanning elements at the cost of optomechanical complexity^53^. Second, out-of-focus fluorescence was visible due to the finite thickness of the light-sheet. As a discriminator for filtering we used the difference in spatial footprint of (almost point-like) neurons and the global large-scale background. To this end, a spatial Fourier-transformation was computed for each 3D volume. A spatial delta-function (neuron) is widely distributed in Fourier space. In contrast, the large-scale fluorescence background and the stripe artifacts are concentrated to distinct regions corresponding to low spatial frequencies. To mask those Fourier regions for filtering, the temporal variance of each pixel in Fourier space was taken (see Methods). After this preprocessing, CaImAn’s motion correction algorithm was applied before temporal 3D segmentation. In the final post-processing step, we applied a volume threshold based on typical neuron morphology. To eliminate false positive neurons caused from drifts in the residual background, the traces were filtered according to their variance. Traces that do not match either of the above criteria were excluded from further analysis. In a final step, the remaining traces were normalized by a linear transformation to the [0,1] interval via *Δ F*/*F* = (*F* − *F*_*min*_)/(*F*_*max*_ − *F*_*min*_). While this normalization scheme homogenizes the amplitude of traces, the information about the number of action potentials is lost. To maintain this sensitivity an alternative normalization was chosen *Δ F*/*F* = (*F* − *F*_*min*_)/ *F*_*min*_ (Suppl. Fig. S5).

**Figure 4.**
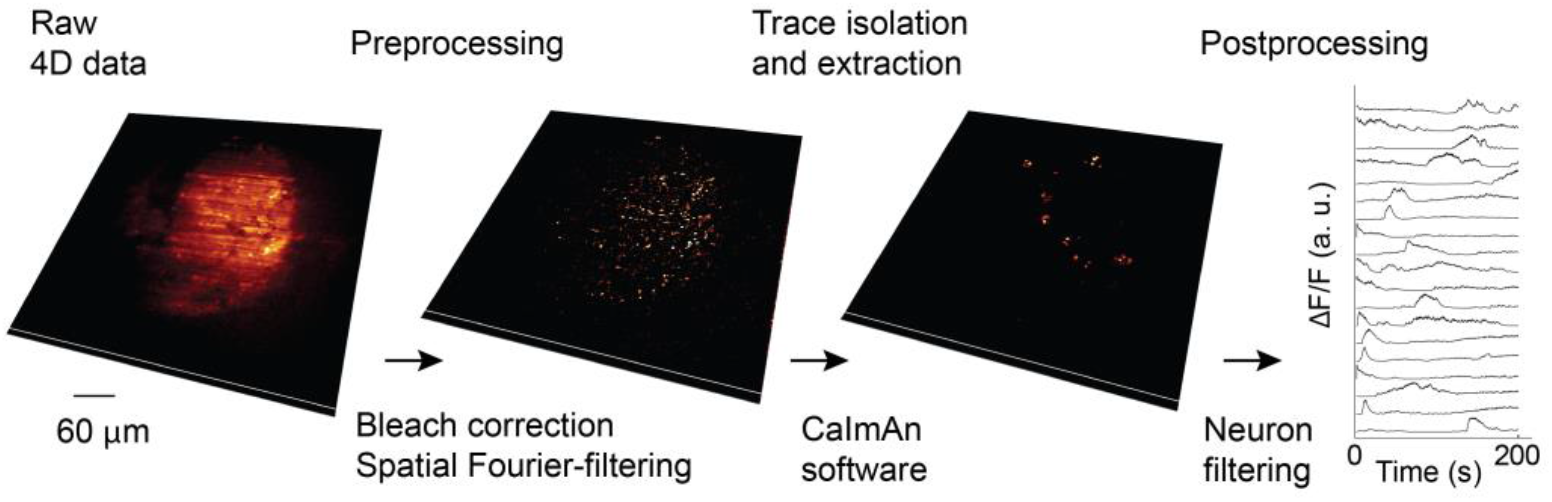
Processing pipeline to extract neuronal signals. To remove the fluorescent background in a first step, the sample is bleach-corrected and spatially filtered via a Fourier filter. Neuronal traces are extracted in 3D using CaImAn^44^. In a final step, neurons are filtered by volume and the temporal traces are filtered according to their variance (see Methods).

## 3. Results

### 3.1 *The 3D analysis* increases *signal-to-noise in activity recordings*

Raw 4D activity recordings displayed fluorescence contributions from active (firing) neurons, a static signal from non-active neurons and out of focus fluorescence due to the microscope’s finite optical sectioning capability (Fig. 5a). As a consequence, the signal-to-noise ratio (SNR) of the raw data is SNR < 1, dominated by the background drift. Using our processing pipeline (Fig. 4) we were able to isolate active neurons from the remaining signal and thus increase the signal to noise ratio by about an order of magnitude. The background-free spatiotemporal Ca^2+^ activity *AC* was obtained and a movie containing only active neurons was distilled (Fig. 5a’). The denoised traces of active neurons were normalized and plotted as a function of time (Fig. 5b, for details see Methods). Comparing with manual annotation on a voxel level, we obtain a F1-score of 0.1 (Suppl. Fig S3, Methods) in the presence of strong background noise. Higher F1-scores can be realized using brighter sensors such as jGCaMP8.

**Figure 5.**
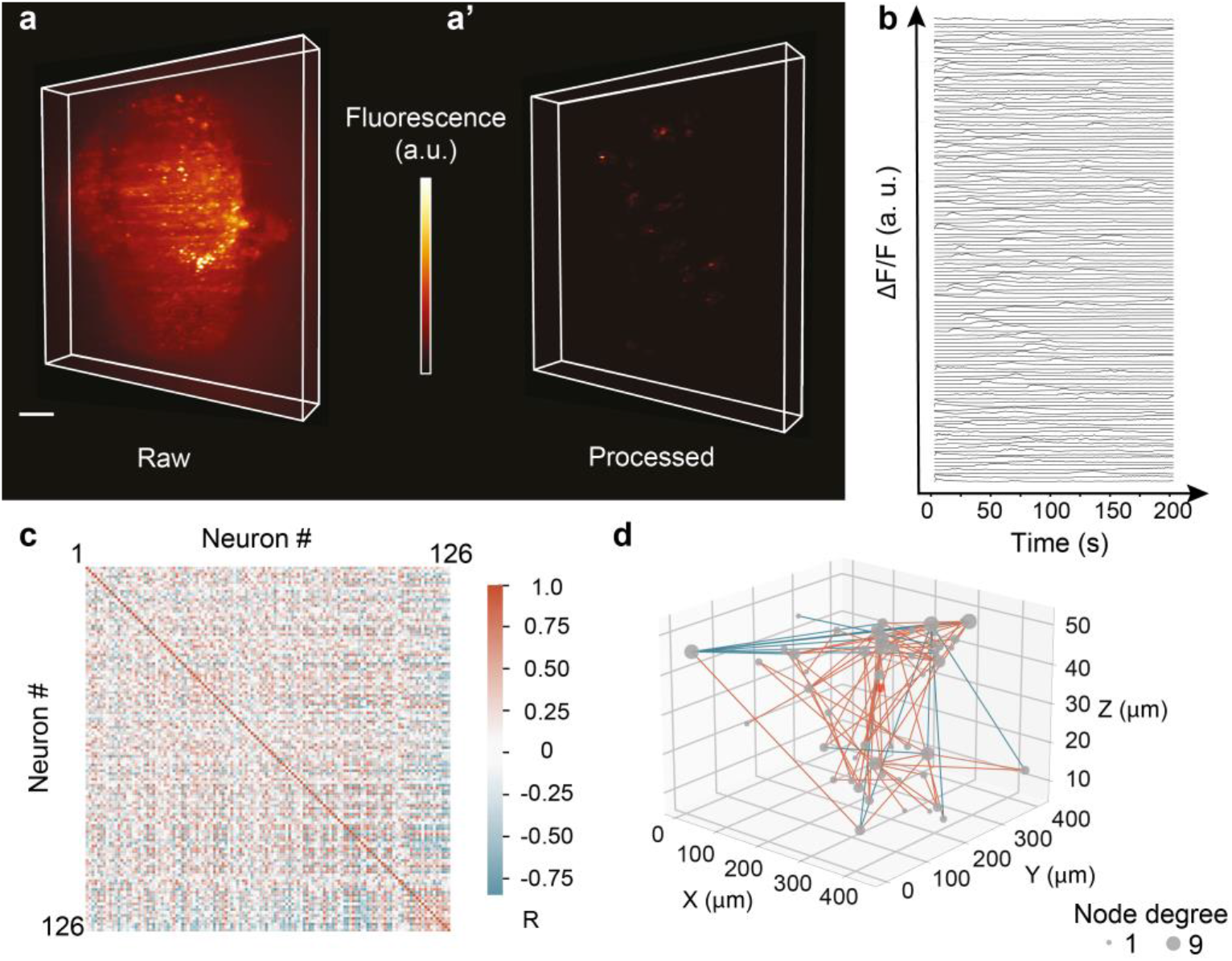
Quantification of neuronal responses in 3D neuron cultures. **(a) 3D lightsheet image of a single timepoint** (R-GECO1.0 calcium sensor, raw data). **(a’)** Same image after the processing pipeline. Only active neurons remain in the observation volume. **(b)** Individually normalized calcium traces. The vertical offset serves display purposes. **(c)** A Spearman rank correlation matrix between any pair of active neurons was constructed using the raw fluorescence traces, ordered according to their distance from the centre of mass (COM). R represents the correlation coefficient value between each pair traces. **(d) 3D functional connectivity map**. Edges are coloured corresponding the R-value of the connection. Only strongest connections with |R|> 0.8 are kept. Nodes’ s size is degree-coded. The red dot represents the COM. Color bar scaling (a.u.): 1584-3778 **(a)**, 0.44-0.64 **(a’)**. Scale bar: 50 µm **(a)**.

### 3.2 3D neuron cultures show patches of correlated activity

To study global connectivity of a network the time-independent pairwise correlation coefficient of two activity traces was obtained^31^ (Fig. 5c) and the matrix storing the respective coefficient values was created. Differences in amplitude of two activity traces can arise from differences in activity or inhomogeneities in illumination. To obtain a correlation measure which is independent of the signal amplitudes but sensitive to common monotonic changes we used the Spearman rank correlation matrix^54^. Neuronal traces were sorted according to their distance from the center of mass (COM) of the neural culture. The correlation matrix (Fig. 5c) revealed distinguished patches of positive and negative correlation formed by neurons located both close to and far from each other. The neuronal network architecture was visualized in 3D using a graph-based representation of the connectivity matrix (Fig. 5d). Neurons were assigned as nodes connected to their neighbors by edges, color-coded using the correlation coefficient value^55^ and thresholded (|R|>0.8)^4^. The final size of the nodes was degree-coded -the more connections a neuron has the bigger the node’s size (Fig. 5d). Interestingly, stronger correlations occurred between adjacent neurons as well as neurons more than 100 *µ*m apart. Further studies using chemical or opto-genetical perturbations could reveal the interplay between far-distant neurons in the network^56^.

## 4. Discussion

We developed a platform for fast volumetric imaging of biological samples by combining a low-complexity custom-built microscope with state-of-the-art computational tools for neuron signal extraction. The setup’s complexity could be kept to a minimum by a plug and play light-sheet module that can be introduced into any standard inverted microscope, including confocal microscopes, where it replaces the condenser. As a proof of principle, we mapped the spontaneous Ca^2+^ activity in the neuronal network of a 3D neuron culture. To show in a proof of principle experiment that Ca-imaging and trace extraction is possible with our setup, neuronal signals have been recorded for 200s in 3D. To test the limits of data acquisition, we recorded beads in agarose gels for as long as 30 min in a continuous stream with the same fast settings we used for Ca-imaging (5Hz/vol, 20 planes/vol) (Suppl. Movie S2). As the full data set fits into the workstation memory (128 GByte), the analysis pipeline is not expected to be a bottleneck. One biophysical limitation is the amount of photo-bleaching, which depends on the laser intensity and the Ca-sensor used. Over the course of the 200s recording (Fig. 5), we observed moderate photo-bleaching using R-GECO1 as a Ca-sensor. Recent sensors such as jGCaMP8 have more than an order of magnitude higher efficiencies, allowing substantially lower laser powers. Therefore, we estimate that recordings with an up to tenfold improvement could be feasible with our setup. Fast, piezo-based mechanical scanning of living samples has been reported before^42^. To minimize mechanical impact on the sample, we kept the load acting on the piezo stage to a minimum (< 100 grams). In addition, we tested actuation of beads embedded in low melting point agarose (Suppl. Movie S2). We find that the lateral sample movement remains at a sub-cellular scale, well in the scope of the built-in motion correction of the analysis pipeline. Spontaneous activity is a hallmark of developing neural circuits where activity patterns along with genetical programs enable their development^57,58^. In particular, synaptic refinement during neural development builds on correlation of pre- and postsynaptic firing^59,60^. The activity correlations we observed for small groups of neurons could initiate the emergence of a functional neuronal circuit^61^. To further investigate this, organoids at other developmental stages could be characterized using our platform. In a recent study by Sharf et al. receivers, senders and brokers nodes were identified in the human brain organoid network using multielectrode arrays^4^. Synchronous network activity at frequencies below 10 Hz (Theta-oscillations) were found. However, the classification of participating cells was challenging due to the limited access for 3D microscopy. Our light-sheet-based approach provides access both to electrophysiology at this timescale as well as neuron type through the imaging of morphological characteristics.

Finally, in order to quantify network formation and function in developing neural organoids, the spiking trains could be deduced, with the ultimate goal to develop and test *in silico* models of the biological neuronal network^62,63^.

## 5. Methods

### 5.1 Light-sheet microscope hardware

The setup features three fiber-coupled diode laser lines (488 nm: Ibeam Smart-488-S-HP, 561 nm: Cobolt Jive, 643 nm: CrystaLaser LC). Light intensity was controlled with a Polychromatic Acousto-Optic Modulator (Neos, PCAOM 48062-2.5-.55). At the fiber exit, the beam gets collimated (F240FC-532, Thorlabs), split on a beam splitter (Thorlabs, BS007) (the generation of two light-sheets is possible) and is further folded by a mirror (Thorlabs, MRA05-E02) and right-angle prisms (Edmund’s Optics, 84-513). Next, the beam hits a cylindrical lens (Edmund’s Optics, 47763) and an aspherical lens (AMS Technologies, 354996A) which results in the formation of a static light-sheet in the field of view. Optical elements were inserted into custom-modified Thorlabs Cage Plates (SP01).

Fluorescence detection is achieved with a water immersion objective (UMPLFLN 20XW Olympus/ 0.5, LUMPLFLN 40XW Olympus/0.8) orthogonally oriented with respect to the illumination. Collected light is detected on a camera (ORCA-Flash4.0 V3 Digital CMOS).

### 5.2 PSF determination

The PSF was obtained by imaging 1% gel-embedded 170 nm fluorescent beads with a 561 nm laser (1% low gelling temperature agarose in ddH_2_O, orange 540/560 nm beads, PS-Speck™ Microscope Point Source Kit P 7220). For 10 beads we measured 4 lateral and 2 axial intensity profiles along any two of the perpendicular axes in three central planes (*xy, xz, yz*) (Fiji) and fitted a Gaussian function

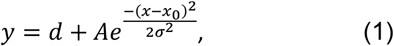

to extract *σ*_*x,y,z*_ as a measure for the lateral and axial resolution.

To estimate the lightsheet thickness, height and homogeneity, the illumination profiles across 200 × 200 µm^2^ of the FOV were mapped. For this purpose, we used a 45° tilted mirror that was moved along the optical axis to deflect the light-sheet directly into the detection lens. The images were fitted with a custom written Python code to extract *σ*_*x*_:

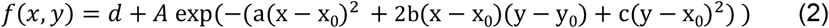

with

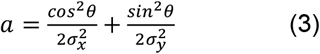

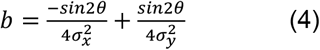

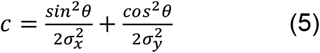

By setting the rotation angle to zero (*θ* = 0) we obtained

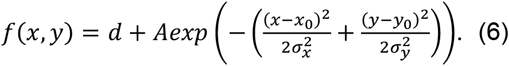

The 1/*e*^2^ beam waist *w*_0_, which corresponds to 2*σ*_*x*_ was plotted as a function of *z*. Data points were fitted with the theoretical description of a Gaussian beam

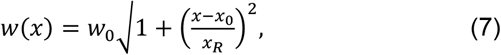

where *x* is the Rayleigh length *x*_0_ corresponds to the shift with respect to the origin.

### 5.3 Microscope control

Computer control was realized with μManager (Suppl. Fig. S1). To attain high acquisition speeds, the camera was set to streaming mode (master). Its clock triggered the Arduino 1, which triggered the remaining hardware (https://micro-manager.org/Arduino). Arduino 2 was controlled using serial commands to provide the number of steps and the direction to the controller of the piezo-positioner (Smaract SDC2). Once armed, the stage waited for triggers from Arduino 1 and moved a predefined distance in closed loop after receiving a trigger. To acquire a time series, the user defines (I) the acquisition parameters in the multi-dimensional acquisition (MDA) tab of μManager (number of frames, exposure time and laser power intensity) and (II) the direction and number of z-planes via the serial command of Arduino 2. For most experiments, the exposure time is set to 10 ms per frame. Scanning was performed in a bidirectional fashion. During acquisition, the light-sheet remained at a pre-set position while the sample was moved stepwise along the z-axis using a piezo-positioner (Smaract, SLC-1730-O20-W-L-E-NM-TI) mounted vertically. Images were saved as separate .tiff files.

### 5.4 Sample preparation and characterization

#### Embryonic stem cell culture (ES)

Mouse embryonic stem cell line (kind gift from Ivan Bedzhov, MPI Münster) were maintained under 5% CO_2_ and 37°C conditions on 10 % gelatine-coated (Sigma, G1393) T25 flasks in DMEM high glucose medium (Sigma D5671) containing 15% FBS (Sigma S0615), 100 U/ml Penicillin-Streptomycin, 2 mM L-Glutamine, 1 mM sodium pyruvate (Sigma, S8636), 0.1 mM NEAA (Sigma, M7145), 0.1 mM 2-mercaptoethanol (Sigma, M3148). Maintenance medium was freshly supplemented with 0.44 nM mLif (Amsbio, AMS-263), 0.4 *µ*M PD 0325901 and 3 *µ*M CHIR 99021 inhibitors (Cayman Chemicals). Seeding cell number was set to 4.2 × 10^5^ cells (for 2-day interval passaging) or 1.5 × 10^5^ (for 3day interval passaging). The cells carry a fluorescent marker in the nucleus (H2B–GFP+) and express a red Ca^2+^ sensor (R-GECO). The cell line was detected positive for myoplasm after the experiments were performed.

#### Induction of neural differentiation of ES cells

The protocol to generate retina organoids was adapted from Völner et al.^45^. In brief, mES cells were dissociated in 0.25% EDTA trypsin (Gibco, 25200056), centrifuged (5min, 1000rpm) and resuspended in differentiation medium to achieve cell count 3 × 10^4^ ml. In order to form embryoid bodies, cells were allowed to reaggregate in U-bottom 96 well plates (3 × 10^3^ cells per well) (Nunc). Differentiation medium was G-MEM-based (Gibco, 21710-025) supplemented with 1 mM sodium pyruvate, 0.1 mM NEAA, 0.1 M 2-mercaptoethanol and 1.5% Knockout Serum Replacement (Gibco, 10828-028). The culture was kept at 5% CO_2_ and 37° C for 7 days, after which each well was filled with additional 200 ul of maturation medium I and transferred to an incubator with 40% O_2_, 5% CO_2_ and 37° C. Maturation medium I was DMEM/F12 supplemented with 100 U/ml Penicillin-Streptomycin and 1% N-2 Supplement (Gibco, 17502048). On day 10, spheroids were di-or tri-sectioned under a stereomicroscope (Nikon, Stereoscope), transferred to low adhesion 24-well plates (Corning, CLS3473) filled with maturation medium II and placed back into the 40% O_2_, 5% CO_2_, 37°C incubator. Maturation medium II was DMEM/F12 freshly supplemented with 100U Pen/Strep, 1% N_2_-suplement, 0.5 *µ*M retinoic acid (ec23®, SRP002), 1 mM taurine (Sigma, T0625). By day 15, the culture was stopped as the sample could not be cultivated further, most likely due to the detected mycoplasma contamination.

#### Cryo-sectioning and immunohistochemistry

We immuno-stained 10 µm-thick sections of the sample obtained with a cryostat (Thermofisher, Cryostar nx70). After imaging, samples were fixed in 4% formaldehyde, incubated overnight in 30% sucrose and immersed in pre-warmed embedding solution (7.5% gelatin and 10% sucrose in DDI water). Blocks containing multiple organoids were cut out and frozen in -50°C isopentane bath. With confocal microscopy (LS980 Zeiss, 40x) we could detect the presence of SOX-2 (ThermoFisher, Invitrogen, eBioscience™, 14-9811-82) and Nestin (Abcam, ab81462) (Fig. 3, Suppl. Fig. S2) within the structures. For TUBB3 (Fig. 3, Suppl. Fig S2) and N-CAD (Suppl. Fig. S2) staining, samples were embedded and cryosectioned directly in Neg-50TM Frozen Section Medium (Thermo Fisher Scientific, 6502B) and stained with anti-N-Cadherin antibody (ThermoFisher, Invitrogen, PA5-29570) and anti-Tubulin ß 3 antibody (Clone Tuj1, BioLegend, MMS-435P) respectively.

### 5.5 Sample mounting

The organoid was placed in an agarose well freshly prepared with the use of a 3D printed mold maker, consisting of two twin plates and a comb. First, the plates were joined together, placed on a clean surface and warm liquid 2% agarose was pipetted between them (2% high gelling temperature in ddH_2_O). The comb was then placed on top of the plates and the mold maker was kept at 4° C. Once the agarose solidified, the comb was lifted and the plates were removed. As a result, we obtained a transparent block of agarose with a hollow well. We cut the well to create a single-use imaging chamber. The chamber was fixed to a bottom glass petri dish between two parafilm stripes using warm liquid agarose. Finally, the organoid was transferred to the well with a cut P200 pipette tip and the dish was filled with medium.

### 5.6 Imaging

Recorded images were binned 4×4 to result in 512×512 pixel at a frame rate of 100 fps. The imaging volume was adjusted to 666 × 666 × 60 μm^3^ with 20 z-planes and the interplane spacing set to 3 *µ*m. We used 10 ms exposure per frame which resulted in a scanning volume rate of 5 Hz. The cells in the organoids expressed a nucleus-localized fusion protein H2B-GFP which allowed us to visualize the tissue’s general architecture. We started by taking a single structural stack using 488 nm laser. Next, we performed functional imaging of the FOV (1000 time points, 200 s) using the 561 nm laser. Raw Ca^2+^ images were re-order and saved as a hyperstack using a custom-written macro (Fiji).

### 5.7 Data processing

#### Preprocessing

To correct for sample bleaching, each frame was normalized by its median intensity:

1. medians_over_time = median_X(Raw_data) #spatial median for each time frame
2. median_normalized = raw_data*1/medians_over_time #for every time frame, normalize by its median
To eliminate lightsheet typical artifacts, including out-of-focus blur and stripe artifacts, a Fourier filter was applied using the following steps:
3. f = FFT_X(Median_normalized) #scipy.fft.rfftn module in python
4. var = VAR_T(f) #stripes and out-of-focus blur show large variance
5. q = 99^th^ QUANTILE(var) #select high variance regions
6. keep_binmask = LOG_10(var) < q #make a mask
7. transform_back = keep_binmask * f #apply mask
8. variance_filtered = IFFT_X(transform_back) #scipy.fft.irfftn in python

#### Trace extraction with CaImAn

To extract spatial footprints of active neurons and their corresponding Ca^2+^ traces we used the CaImAn 1.9.9 package. First, the motion correction was performed. Then, CNMF algorithm was run twice on the motion corrected and memory mapped data. For motion correction, relevant parameters were set to

- strides = (18, 18, 4), overlaps = (9, 9, 2), max_shifts = (4, 4, 2), max_deviation_rigid = 5

For the CNMF fit

- rf = 10, stride = 10, K = 10, gSig = [2,2,2], merge_thresh = 0.8, nb = 2, p = 2

were selected. After the first fit, identified components were evaluated and only those displaying a peak SNR ≥ 1 and a spatial correlation ≥ 0.5 were accepted. The CNMF was then run again on the whole volume seeded with the accepted components. Each denoised trace was individually normalized *Δ F*/*F = (F* − *F*_min_*)/*(*F*_max_ − *F*_min_), where *F*_max_ and *F*_min_ were the highest and lowest intensities recorded in the trace, respectively. Spearman rank correlation matrix was constructed using fluorescent traces of active neurons, ordered according to their distance from the organoid’s center of mass using the seaborn package (0.11.2). Using the spatial coordinates of identified components, we could reconstruct the network as a graph, where nodes represent neurons and edges the interactions between them. The edges adapt the colors corresponding to the R-value of the connection as calculated previously in the matrix. To display only the strongest interactions, a threshold was set on edges to show connections with |R|> 0.8. The node size is then degree-coded so potential hubs in the network can be identified.

#### Postprocessing

We chose as spatial thresholds a minimum neuron volume of 710 µm^3^ and a maximum volume of 7605 µm^3^, estimated from manual annotations of the raw data. The variance threshold for the filtering of the fluorescence traces was chosen empirically to 0.06 after normalizing the traces to [0,1] interval.

#### CaImAn segmentation performance

To estimate the performance of our software pipeline we annotated 3 regions from 3 slices in the dataset from figure 5 (z-slice=6,12,18) by hand. Three annotators independently annotated the slices, and then the intersection of the annotations was taken as a “consensus annotation”. We overlaid the consensus annotation to the spatial segmentation of the CaImAn software and calculate precision, recall and F1-score metrics. We obtain a precision of 0.057, a recall of 0.424 and an F1-score of 0.100 when comparing the consensus annotation with the CaImAn result (Suppl. Figure S3). This evaluates the spatial segmentation performance on a single voxel level, which is a rather strict quality criterion. Also, since the annotations are performed on the raw data, which is not motion corrected yet, motion artifacts systematically lower the performance metrics. Using brighter Ca-sensors (e.g. GCaMP7) can significantly increase segmentation performance due to an expected improvement in SNR>10.

## Acknowledgments

We thank Otger Campas and Paul Hansma for initial discussions, Carolyne Sheline and Nicoleta Dahnovici for contributions in the early phases of the project, Jacopo Di Russo and Jennifer Diemer for help with the stem cell culture and Joachim Spatz for his generous support. In addition, we thank Johannes Friedrich and Andrea Giovannuci for stimulating discussions.

## Competing interests

The authors declare no conflicts of interest.

## Data availability

Data underlying the results presented in this paper are not publicly available at this time but may be obtained from Friedhelm Serwane upon request.

## Funding information

This project has received funding from the European Research Council (ERC) under the European Union’s Horizon 2020 research and innovation program (Grant Agreement No. 850691), the Center for NanoScience (CeNS) at the Ludwig-Maximilian-University (LMU) Munich, and the Munich Cluster for Systems Neurology (SyNergy). P.M.W. and F.S. acknowledge support from the Baden-Württemberg Stiftung (MIVT-5).

## Author contributions

P.W. characterized the experimental setup and performed organoid culture experiments. P.W., F.K. and E.S. analysed the data, K.S. and S.S. performed cryo-sectioning and immuno-fluorescence imaging, F.S. designed research and supervised the project. P.W. and F.S. wrote the paper. All authors reviewed the manuscript.

## Notes

### Competing Interest Statement

The authors have declared no competing interest.

### Summary of Updates

The clarity of the main text has been improved. The figures have been updated with respect to readability.

