## Supplementary Information for "A minimal-complexity light-sheet microscope maps network activity in 3D neuronal systems"

Suppl. Table 1: Existing Ca-imaging setups. Few, low complexity setups exist as addons to standard inverted microscopes which offer volumetric readout at single neuron resolution.

| **Type** | **FOV (μm³)** | **Rate (Hz)** | **Single-neuron resolution in 3D** | **Complexity** | **Reference** |
|---|---|---|---|---|---|
| **LS** | 665 x 665 x 60 | 5 | y | add-on | This work |
| **LMF** | 350 x 350 x 30 | 5 | y | add-on | Prevedel et al. 2014[1] |
| **LS** | 3000×3000×7000 | NA | y | stand alone | Liu et al., 2021[2] |
| **LS** | NA | NA | y | add-on | Bruns et al., 2016[3] |
| **LS** | 800 x 600 x 200 | 0.8 | y | stand alone | Ahrens et al., 2013[4] |
| **LS** | 100 x 800 x 40 | 5 | y | stand alone | Panier et al., 2013[5] |
| **LS** | 830 x 430 x 200 | 2-3 | y | stand alone | Vladimirov et al., 2014[6] |
| **1P and 2P LS** | 500 x 200 x 200 | 5 | y | stand alone | Lemon et al., 2015[7] |
| **2P with TeFo** | 500 x 500 x 500 | 5.7 | n | stand alone | Prevedel et al., 2016[8] |
| **LFM with EDoF** | 416 x 832 x 160 | 33 | NA | stand alone | Quirin et al., 2016[9] |
| **SCAPE** | 600 × 650 × 134 | 10 | y | stand alone | Bouchard et al., 2015[10] |
| **SCAPE** | 392 × 299 × 41 | 25.75 | y | stand alone | Voleti et al., 2019[11] |
| **LFM AI-enhanced** | 350 x 280 x 120 | 10 | y | stand alone | Wagner et al.,2021[12] |
| **OCPI** | 223 x 127 x 200 | 20 | y | stand alone | Greer et al., 2018[13] |
| **confocal LFM** | ∅ 800 x 200 | 6 | y | stand alone | Zhang et al., 2020[14] |
| **2P with ETL** | 500 x 500 x 100 | 4 | n | stand alone | dal Maschio et al., 2017[15] |
| **2P with ETL** | 500 x 500 x 450 | 10 | n | stand alone | Han et al., 2019[16] |
| **HyMS with TeFo** | 765 x 665 x 800 | 4.3-13 | n/y | stand alone | Weisenburger et al., 2019[17] |
| **TPM with OPLUL** | 375 x 112 x 130 | 14 | n | stand alone | Kong et al, 2015.[18] |
| **LS** | NA | 1.5 | n | stand alone | Markov et al., 2019[19] |
| **LBM** | 600 x 600 x 500 | 10 | n | stand alone | Demas et al., 2021[20] |
| **miniaturized 2P** | 510 x 510 x 40 | 7.5 | n | stand alone | Zong et al., 2021[21] |
| **OPM** | 500 x 300 x 200 | 3.3 | y | stand alone | Yang et al.2022[22] |

2PLSM = 2-Photon laser scanning microscopy
CADoF = Continuously Adjustable Depth of Focus
EDoF = Extended Depth of Field
ETL = Electrically Tunable Lens
HyMS = Hybrid Multiplexed Sculpted Light Microscopy
LBM = Light Beads Microscopy
LS = Light-sheet
LFM = Light Field Microscope
SCAPE = Swept Confocally-Aligned Planar Excitation
STM = Scanning Two-photon Microscopy
TeFo = Temporal Focusing
OCPI = Objective Coupled Planar Illumination
OPLUL = Optical Phase-Locked Ultrasound Lens
OPM = Oblique Plane Microscopy

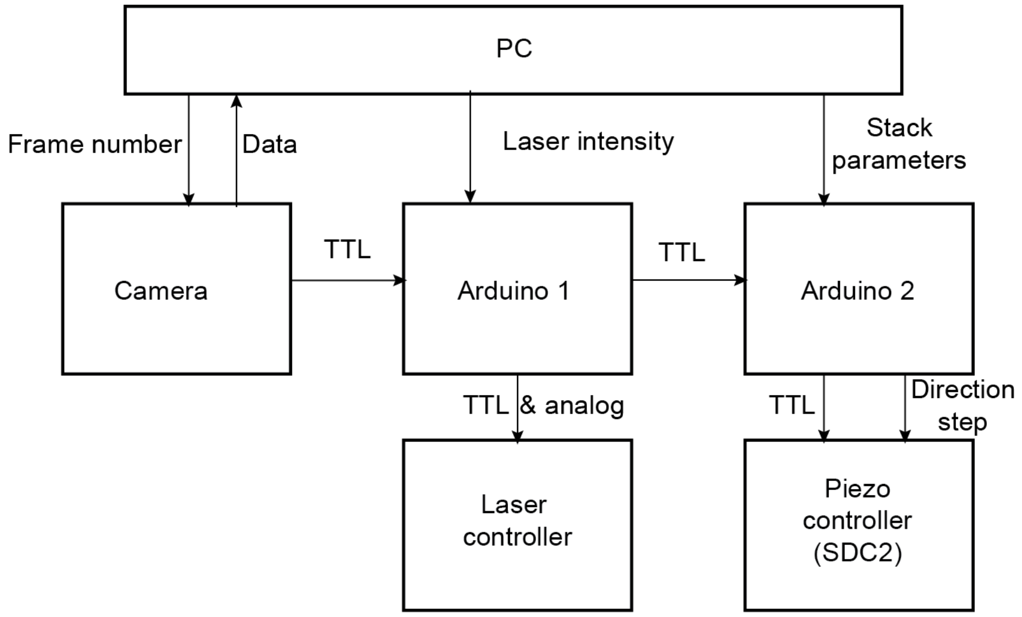


Supplementary Figure S1: Electronic control of the light-sheet microscope. The PC sets the number of frames, the laser intensity and the stack parameters. The camera acts as the master trigger to control Arduino 1. This, in turn sets the laser power and triggers the z-stage Arduino 2.

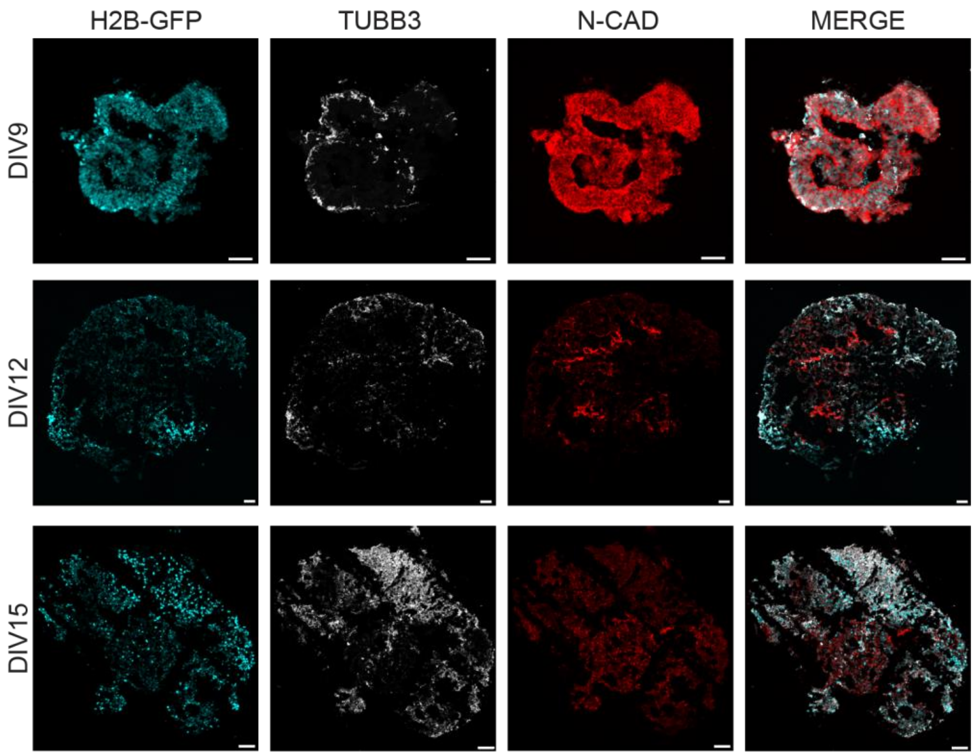


Supplementary Figure S2: Immunofluorescence images of cryo-sectioned organoids on DIV 9, 12 and 15. TUBB3 indicates neuronal identity and is most prominent on the outer layers of the organoid which were used for calcium signal analysis. Scale bar: 50 μm.

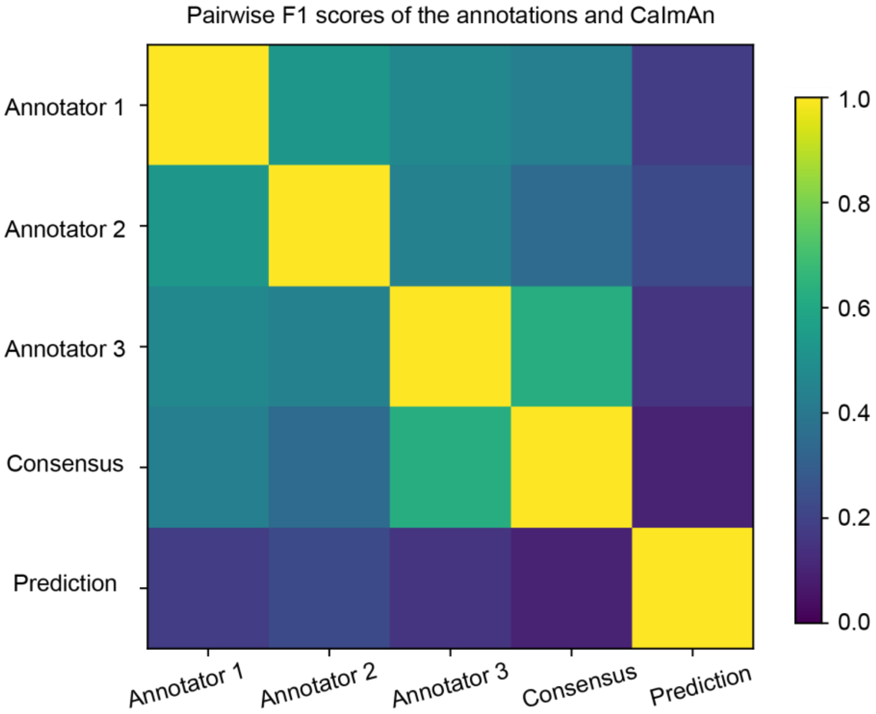


Supplementary Figure S3: Evaluation of the segmentation performance. The annotations from 3 human annotators and consensus are numerically compared against the segmentation by our software pipeline. All annotations are compared to each other, including the segmentation from our pipeline, using the F1-score.

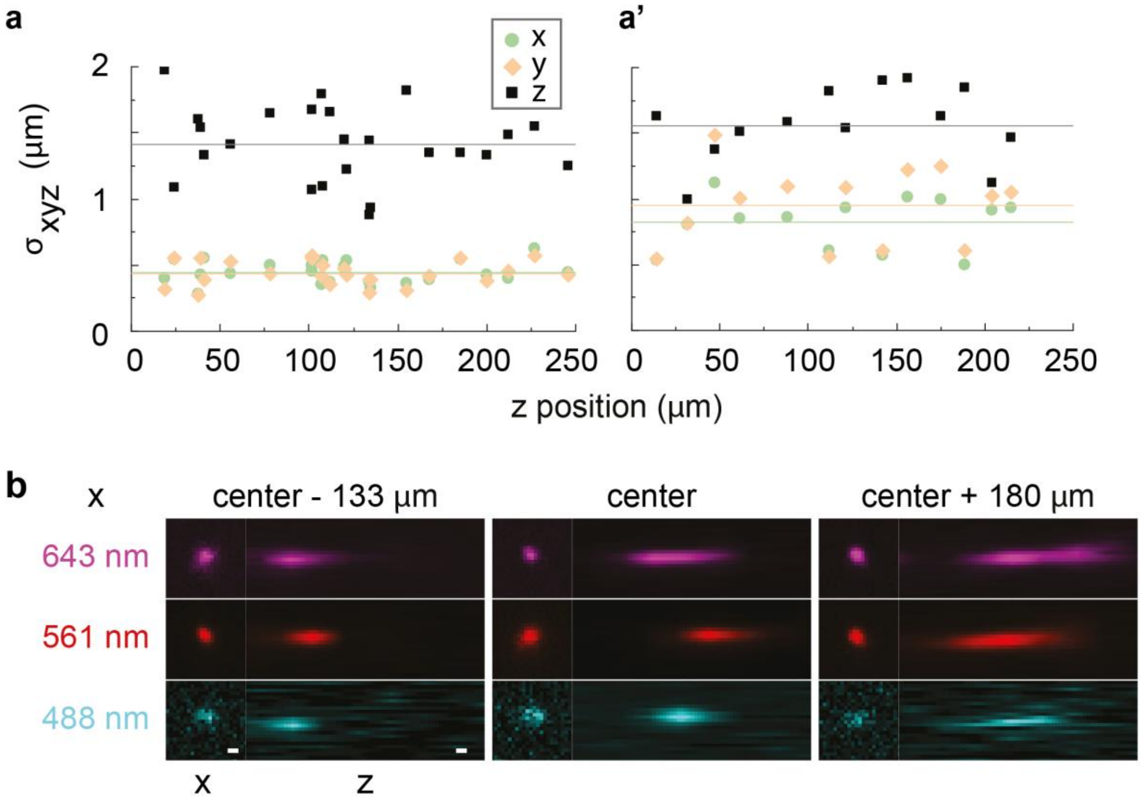


Supplementary Figure S4: PSF characterization of 200nm beads embedded in 1% low melting point agarose. (**a,a'**) Resolution as a function of imaging depth using the water dipping objectives in immersion mode (a: 40x, a': 20x). The resolution is not degraded for increasing z-positions. (**b**) Resolution for simultaneous multicolor illumination at 3 exemplary x-positions. While multicolor imaging is possible, the axial resolution varies as a function of x. Solid lines indicate the mean. Scale bar: 1 $\mu$m (b).

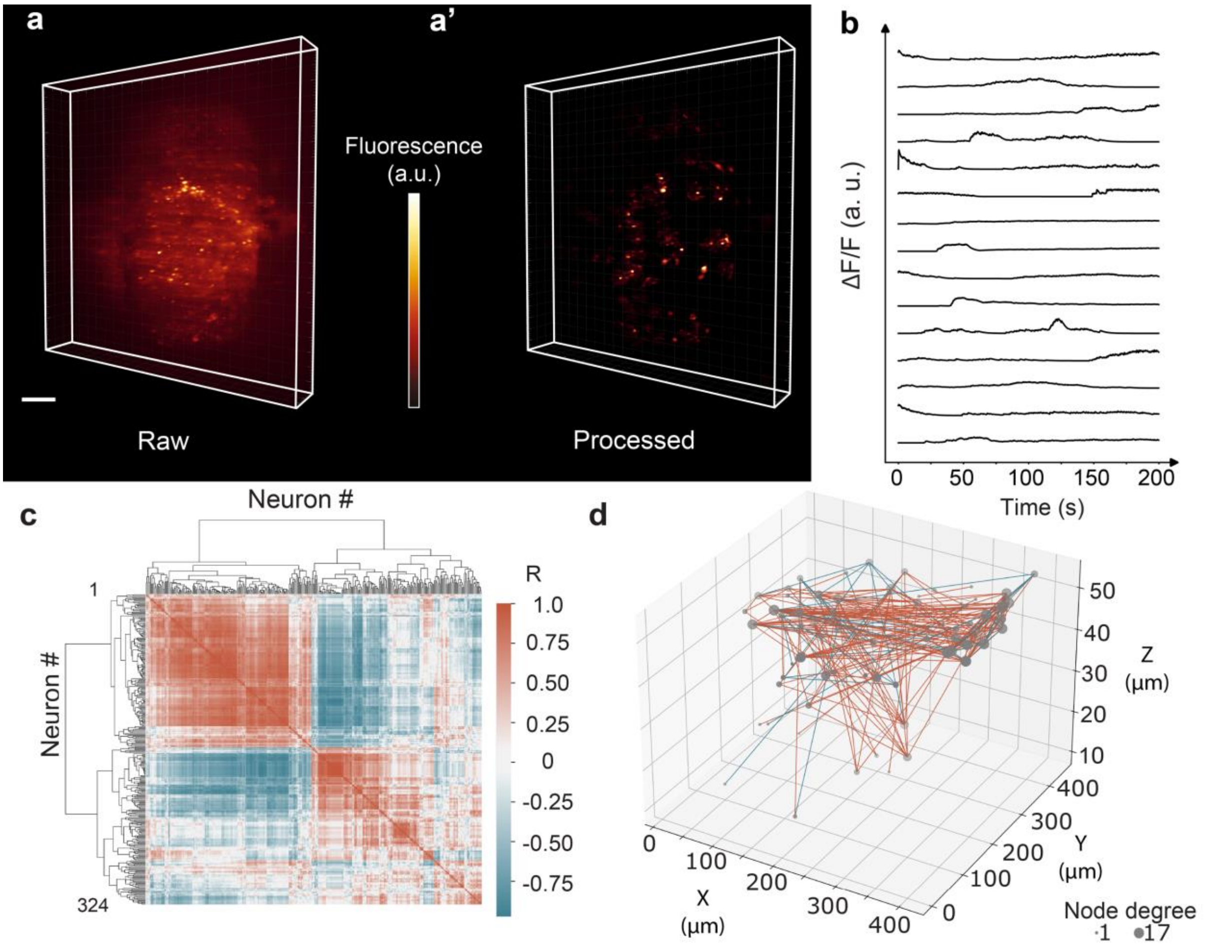


Supplementary Figure S5: Data of Fig. 5 with trace normalization $\Delta F/F = (F - F_{min})/F_{min}$ and hierarchical clustering. (**a,a'**) Raw and processed 3D lightsheet image of a single timepoint (R-GECO1.0 calcium sensor). (**b**) Representative calcium traces. (**c**) Spearman rank correlation matrix using the unweighted pair group method with arithmetic mean, UPGMA, https://en.wikipedia.org/wiki/UPGMA. More traces are found due to the different normalization and the absence of the variance filter (see main text) (**d**) 3D functional connectivity map. Edges are colored corresponding the R-value of the connection (**c**). Only strongest connections with |R|> 0.95 are kept. The node size is degree-coded. Color bar scaling (a.u.): 1584- 4888.54 (a), 0.47-0.63 (a'). Scale bar: 50 $\mu$m (a).

Supplementary Movie 1: Side by side comparison of raw and processed data of Figure 5.

Supplementary Movie 2: Maximum intensity projection of a 30 min timelapse of 200 nm beads embedded in 1% low melting point agarose. The movie was recorded using the same acquisition settings as the data obtained in Fig. 5 (5 Hz/volume, 20 planes/volume).

Bibiography


1 Prevedel, R. *et al.* Simultaneous whole-animal 3D imaging of neuronal activity using light- field microscopy. *Nature Methods* **11**, 727–730, doi:10.1038/nmeth.2964 (2014).

2 Liu, Y., Rollins, A. M. & Jenkins, M. W. CompassLSM: axially swept light-sheet microscopy made simple. *Biomed. Opt. Express* **12**, 6571-6589, doi:10.1364/BOE.440292 (2021).

3 Bruns, T. *et al.* Miniaturized modules for light sheet microscopy with low chromatic aberration. *Journal of Microscopy* **264**, 261-267, doi:https://doi.org/10.1111/jmi.12439 (2016).

4 Ahrens, M. B., Orger, M. B., Robson, D. N., Li, J. M. & Keller, P. J. Whole-brain functional imaging at cellular resolution using light-sheet microscopy. *Nature Methods* **10**, 413-420, doi:10.1038/nmeth.2434 (2013).

5 Panier, T. *et al.* Fast functional imaging of multiple brain regions in intact zebrafish larvae using Selective Plane Illumination Microscopy. *Frontiers in Neural Circuits* **7**, 65, doi:10.3389/fncir.2013.00065 (2013).

6 Vladimirov, N. *et al.* Light-sheet functional imaging in fictively behaving zebrafish. *Nature Methods* **11**, 883–884, doi:10.1038/nmeth.3040 (2014).

7 Lemon, W. C. *et al.* Whole-central nervous system functional imaging in larval Drosophila. *Nature Communications* **6**, 1-16, doi:10.1038/ncomms8924 (2015).

8 Prevedel, R. *et al.* Fast volumetric calcium imaging across multiple cortical layers using sculpted light. *Nature Methods* **13**, 1021–1028 doi:10.1038/nmeth.4040 (2016).

9 Quirin, S. *et al.* Calcium imaging of neural circuits with extended depth-of-field light-sheet microscopy. *Opt Lett* **41**, 855-858, doi:10.1364/OL.41.000855 (2016).

10 Bouchard, M. B. *et al.* Swept confocally-aligned planar excitation (SCAPE) microscopy for high speed volumetric imaging of behaving organisms. *Nature Photonics* **9**, 113-119, doi:10.1038/nphoton.2014.323 (2015).

11 Voleti, V. *et al.* Real-time volumetric microscopy of in vivo dynamics and large-scale samples with SCAPE 2.0. *Nature Methods* **16**, 1054-1062, doi:10.1038/s41592-019-0579-4 (2019).

12 Wagner, N. *et al.* Deep learning-enhanced light-field imaging with continuous validation. *Nature Methods* **18**, 557–563, doi:10.1038/s41592-021-01136-0 (2021).

13 Greer, C. J. & Holy, T. E. Fast objective coupled planar illumination microscopy. *Nature Communications* **10**, doi:10.1038/s41467-019-12340-0 (2019).

14 Zhang, Z. *et al.* Imaging volumetric dynamics at high speed in mouse and zebrafish brain with confocal light field microscopy. *Nature Biotechnology* **39**, 74–83, doi:10.1038/s41587-020-0628-7 (2020).

15 dal Maschio, M., Donovan, J. C., Helmbrecht, T. O. & Baier, H. Linking Neurons to Network Function and Behavior by Two-Photon Holographic Optogenetics and Volumetric Imaging. *Neuron* **94**, 774-789, doi:10.1016/j.neuron.2017.04.034 (2017).

16 Han, S., Yang, W. & Yuste, R. Two-Color Volumetric Imaging of Neuronal Activity of Cortical Columns. *Cell Reports* **27**, 2229-2240, doi:10.1016/j.celrep.2019.04.075 (2019).

17 Weisenburger, S. *et al.* Volumetric Ca2+ Imaging in the Mouse Brain Using Hybrid Multiplexed Sculpted Light Microscopy. *Cell* **177**, 1050-1066, doi:10.1016/j.cell.2019.03.011 (2019).

18 Kong, L. *et al.* Continuous volumetric imaging via an optical phase-locked ultrasound lens. *Nat Methods* **12**, 759-762, doi:10.1038/nmeth.3476 (2015).

19 Markov, D. A., Petrucco, L., Kist, A. M. & Portugues, R. A cerebellar internal model calibrates a feedback controller involved in sensorimotor control. *Nature Communications* **12**, 1-21, doi:10.1038/s41467-021-26988-0 (2021).

20 Demas, J. *et al.* High-speed, cortex-wide volumetric recording of neuroactivity at cellular resolution using light beads microscopy. *Nature Methods* **18**, 1103-1111, doi:10.1038/s41592-021-01239-8 (2021).

21 Zong, W. *et al.* Large-scale two-photon calcium imaging in freely moving mice. *Cell* **185**, 1240-1256, doi:10.1101/2021.09.20.461015 (2021).

22 Yang, B. *et al.* DaXi—high-resolution, large imaging volume and multi-view single-objective light-sheet microscopy. *Nature Methods* **19**, 461-469, doi:10.1038/s41592-022-01417-2 (2022).
